# Oral thermosensing by murine trigeminal neurons: modulation by capsaicin, menthol, and mustard oil

**DOI:** 10.1101/486480

**Authors:** Sara C.M. Leijon, Amanda F. Neves, Joseph M. Breza, Sidney A. Simon, Nirupa Chaudhari, Stephen D. Roper

## Abstract

When consumed with foods, mint, mustard and chili peppers generate pronounced oral thermosensations. Here we recorded responses in mouse trigeminal ganglion neurons to investigate interactions between thermal sensing and the active ingredients of these plants — menthol, allyl isothiocyanate (AITC), and capsaicin, respectively — at concentrations found in foods and commercial hygiene products. We carried out *in vivo* confocal calcium imaging of trigeminal ganglia in which neurons express GCaMP3 or GCAMP6s and recorded their responses to oral stimulation with thermal and the above chemesthetic stimuli. In the V3 (oral sensory) region of the ganglion, thermoreceptive neurons accounted for ∼10% of imaged neurons. We categorized them into 3 distinct classes: cool-responsive and warm-responsive thermosensors, and nociceptors (responsive only to temperatures ≥43-45°). Menthol, AITC, and capsaicin also elicited robust calcium responses that differed markedly in their latencies and durations. Most of the neurons that responded to these chemesthetic stimuli were also thermosensitive. Capsaicin and AITC increased the numbers of warm-responding neurons and shifted the nociceptor threshold to lower temperatures. Menthol attenuated the responses in all classes of thermoreceptors. Our data show that while individual neurons may respond to a narrow temperature range (or even bimodally), taken collectively, the population is able to report on graded changes of temperature. Our findings also substantiate an explanation for the thermal sensations experienced when one consumes pungent spices or mint.

**Key points summary:**

- Orosensory thermal trigeminal afferent neurons respond to cool, warm, and nociceptive hot temperatures with the majority activated in the cool range.
- Many of these thermosensitive trigeminal orosensory afferent neurons also respond to capsaicin, menthol and/or mustard oil (allyl isothiocyanate, AITC) at concentrations found in foods and spices.
- There is significant but incomplete overlap between afferent trigeminal neurons that respond to heat and to the above chemesthetic compounds.
- Capsaicin sensitizes warm trigeminal thermoreceptors and orosensory nociceptors; menthol attenuates cool thermoresponses.

## Introduction

Somatosensory afferent fibers in the tongue, palate, and buccal cavity are activated by a variety of stimuli, including mechanosensory, nociceptive, thermal, and chemesthetic. Moreover, chemesthetic stimuli from naturally occurring compounds found in foods, beverages, spices, and oral hygiene products significantly affect other somatosensations. For instance, capsaicin, the active component of hot chili peppers, evokes the perception of heat and pain and sensitizes the oral mucosa to subsequent warm stimuli (Green, 2005). These effects are attributed in large part to capsaicin acting on TRPV1 channels expressed in afferent thermosensitive nerve fibers that innervate the oral mucosa (Roper, 2014; Simon and Gutierrez, 2017). Similarly, mustard oil (allyl isothiocyanate, AITC), the active ingredient in wasabi, horseradish, and mustard, causes an “irritating” pungent sensation (Albin et al., 2008). These sensations might be explained by AITC stimulating TRPV1 and TRPA1 channels in primary sensory afferents located there. By contrast, menthol, found in mint, elicits coolness in the mouth (Green, 1985), likely by activating TRPM8-expressing sensory afferent fibers (Bautista et al., 2007).

Thermal sensations in the oral cavity are ascribed to activation of somatosensory (trigeminal) afferent fibers, although gustatory afferent nerves and sensory neurons in the geniculate ganglion have long been known also to respond to thermal stimuli (Zotterman, 1935; Zotterman, 1936; Fishman, 1957; Lundy & Contreras, 1999; Yokota & Bradley, 2016, 2017). Interest in oral thermoreception dates back to the earliest studies obtained with single fiber recording methodologies (Zotterman, 1935). However, details regarding the different types of thermoreceptors that innervate the oral mucosa and their relationship to chemesthetic stimuli were lacking. Recently, functional imaging of afferent ganglion neuronal activity has begun to answer those questions (Yarmolinsky et al., 2016), but left a gap regarding how the active ingredients of natural spices, applied at concentrations encountered in foods and commercial products, stimulate thermal sensations. To address this, we have undertaken a study using scanning laser confocal calcium imaging of trigeminal ganglion sensory neurons in anesthetized mice during oral thermostimulation. Moreover, we have investigated how thermosensations are modified by oral lavage with concentrations of capsaicin, AITC, and menthol that are found in foods and spices and that elicit chemesthetic sensations (pungency, irritation, cooling) in human subjects. We have quantified and resolved the cellular basis for the interaction between these compounds and thermal sensing. The findings indicate that, at least in mice, trigeminal neurons responding to oral cooling far outnumber those that respond to warming, and that chemesthetic compounds have distinct and significant effects on the oral thermoreceptors. Namely, capsaicin and AITC sensitize nociceptors and warm-responding neurons and menthol attenuates responses mainly in cool thermoreceptors. Our data are also consistent with the notion that neural coding of oral thermosensations can be explained by population coding, as postulated for cutaneous thermosensitivity (Ma, 2012; Wang et al., 2018). Abstracts of these findings have been published (Leijon et al., 2016; Leijon SCM et al., 2017, 2018).

## Materials & Methods

### Animals

We used mice that express GCaMP3 as a knock-in/knockout at the Pirt locus (obtained from X. Dong, Johns Hopkins). These mice have previously been reported to express GCaMP in all sensory neurons of the dorsal root, trigeminal, and geniculate ganglia (Kim et al., 2014; Wu et al., 2015; Dvoryanchikov et al., 2017). In addition, we used GCaMP6s-expressing mice generated by crossing floxed GCaMP6s mice (Jax stock #024106) with Pirt-Cre mice (obtained from X. Dong, Johns Hopkins). All mice were backcrossed to C57Bl/6 for 8-10 generations. Adult mice (3-8 months old) of both sexes were used. We observed no difference in the results between male versus female mice or GCaMP3 versus GCaMP6s mice, although GCaMP6s fluorescent signals were stronger. Data included in this study were obtained from 23 animals. Mice were housed with a 12-hour light cycle with *ad libitum* access to food and water. Experiments were conducted during daytime in the research laboratory. All procedures for surgery and euthanasia were reviewed and approved by the University of Miami IACUC (approval reference 16-208). The euthanasia was performed by CO_2_ asphyxiation followed by cervical dislocation as secondary method of euthanasia.

### Reagents

All reagents were purchased from Sigma. Capsaicin (12084), L-menthol (W266523), and mustard oil (allyl isothiocyanate, AITC; 377430) were dissolved in 100 % DMSO, and diluted to their final concentrations in artificial saliva (15 mM NaCl, 22 mM KCl, 3 mM CaCl_2_, 0.6 mM MgCl_2_, pH 5.8). The DMSO concentrations in the final solutions were 0.06-0.2% for capsaicin, 2.0% for AITC, and 3% for menthol. We confirmed in control experiments that 3% DMSO in artificial saliva does not evoke responses in trigeminal neurons, nor does it affect temperature responses. For rinsing the exposed trigeminal ganglion, we used Tyrode’s buffer (140 mM NaCl, 5 mM KCl, 1 mM MgCl_2_, 2 mM CaCl_2_, 9.4 mM HEPES, 10 mM glucose, 10 mM Na pyruvate, 5 mM NaHCO_3_, pH 7.2-7.4).

### Surgical preparation

Mice were anesthetized with ketamine and xylazine (intraperitoneally 120 mg/kg ketamine, 10 mg/kg xylazine) and placed supine on a far infrared surgical warming pad (DCT-15, Kent Scientific). The animal’s core temperature was monitored with a rectal probe and maintained between 35 and 36 °C. Depth of anesthesia was monitored by the hind paw withdrawal reflex; ketamine booster injections (intraperitoneal) were administered to ensure continued surgical plane of anesthesia throughout the surgery and imaging session. Animals never recovered from anesthesia during or after the procedures (terminal surgery) or during imaging, and mice were euthanized by CO_2_ asphyxiation followed by cervical dislocation immediately after the functional imaging.

To facilitate respiration during oral stimulus presentation, the trachea was exposed and cannulated. A flexible tube was passed through the esophagus to produce a uniform stimulus delivery into the oral cavity (Sollars & Hill, 2005; Wu et al., 2015). For surgical exposure of the region of the trigeminal ganglion that innervates the oral cavity, V3, mice were placed in prone position. Muscle tissue on the dorsolateral side of the skull was cauterized, the zygomatic arch removed, and a small cranial window was opened. A partial hemispherectomy was performed by careful aspiration to allow optical access to the trigeminal ganglion. The head was stabilized by affixing a custom head holder to the skull with a nylon screw and dental acrylic. From the moment of exposure, the ganglion was intermittently rinsed with 35 °C Tyrode’s solution to maintain a favorable neuronal environment.

### Functional imaging and data collection

The surgically prepared mouse was transferred to the stage of an Olympus FV1000 confocal microscope equipped with a 10x-long working distance objective, (UPlanFl, N.A. 0.3). Confocal scans of GCaMP3-/GCAMP6s-labeled ganglion neurons using 488-nm laser excitation with a 505–605 nm emitter filter were taken at ∼1 Hz, and an area of approximately 1,000,000 square microns (640 x 400 pixel size) was covered. For thermal stimulation, we applied cooled or heated artificial saliva (5 ml over 10 sec) at 90 sec intervals. Solutions of artificial saliva at ∼0 °C (ice-cooled) and ∼55 °C were mixed in different proportions to achieve the specified temperatures in the oral cavity. Between thermal stimuli, the oral temperature was returned to resting oral temperature by continuous flow of artificial saliva (∼4 ml/min) at 32 °C. Oral temperature was continuously monitored and recorded by a probe inserted in the buccal cavity and connected to a wireless transmitter (UWBT-TC-UST-NA, OMEGA Engineering, Inc.). Baseline fluorescence was recorded for a minimum of 30-60 sec at the beginning of each recording. When testing the interactions of the chemesthetic compounds– capsaicin, menthol, and AITC–on temperature, two replicates of temperature-response relations were recorded (controls), followed by application of the chemesthetic compound at 5 ml over 10 sec. Following this, we carried out a third repeat of the temperature-response series. In separate experiments when testing latency and duration, the chemesthetic compounds were applied at 1 ml over 10 sec without a subsequent continuous flow of artificial saliva. In trials where all three compounds were tested sequentially, their order of application was counterbalanced between experiments. The compounds were diluted in artificial saliva to the following final concentrations: capsaicin 30-100 µM; menthol, 3 mM; and AITC, 10 mM, consistent with their concentrations in foods and commercial hygiene products (see below).

### Data analysis and statistics

Scans were recorded with Fluoview v. 2 software (Olympus) and digitized/stabilized using FIJI (ImageJ). Baseline-subtracted scans were analyzed to identify responding neurons. To minimize subject bias, regions of interest (ROIs) were manually drawn over any neuron showing fluorescence changes, regardless of stimulus application. ROIs were subsequently analyzed using MATLAB (v. R2017b). We used custom MATLAB code, modified from Wu et al. (2015). The modification consisted of an additional part in the script that corrects for baseline drift. The full code is available upon request. Responses (ΔCa2+_i_) were quantified as peak fluorescence change, ΔF, divided by baseline fluorescence, F_0_ (Δ*F*/*F*_0_). The acceptance criterion for responses was Δ*F*/*F*_0_ > 5 s.d. baseline fluorescence. For thermal stimuli, acceptance criteria also included that responses occurred at consistent latencies after stimulus onset. Neurons that stopped responding in subsequent recordings (drop-outs) were excluded from the analyses.

By collapsing the slices in a stack of scans (i.e., creating a projection), we obtained excellent resolution of the imaged trigeminal neurons and could count the total number of neurons in the field. This allowed us to calculate the incidence and measure the diameters of responding neurons. Cell sizes were only measured in neurons with visible nuclei.

Sample sizes are indicated in main text or figure legends. In the text, samples are presented as mean ± s.d. In figures, error bars show standard deviations. Unless otherwise specified, statistical significance was determined using two-tailed Student’s t-test between two groups, otherwise by unpaired one-way ANOVA, using Prism v.6 (GraphPad). For temperature response relations, the same neurons were monitored throughout the pre- and post-conditions, allowing for paired tests. Statistical significance was defined as p<0.05.

## Results

When the oral cavity in mice was flushed with artificial saliva at temperatures ranging from 5° to 53°, sensory neurons in the V3 region of the trigeminal ganglion (TG) showed robust responses (Fig. 1). About 10% of trigeminal neurons in the region that was imaged were thermosensitive (1090/11,700 neurons; 10 mice). Thermosensitive trigeminal neurons responded to oral lavage with artificial saliva at temperatures above and below the resting temperature in the mouse oral cavity (32 °C). We found that thermally sensitive TG neurons that innervate the oral mucosa fell into 3 ranges: (1) cool-responding neurons that were activated by temperatures <32°; (2) warm-responding neurons, activated >32°; and (3) a group of neurons that responded to stimuli only > 43°-45°. Given this high temperature threshold that is consistent with cutaneous nociceptors (Hardy et al., 1951), we define this third group as nociceptors. A population of neurons exhibited bimodal thermosensitivity, that is, they responded to cool and warm alike. We did not find any obvious spatial mapping of the different thermosensitive groups in the V3 area of the trigeminal ganglion (Fig. 1a). Our data confirm and, as will be shown below, extend the findings of Yarmolinsky et al. (2016).

**Figure 1.**
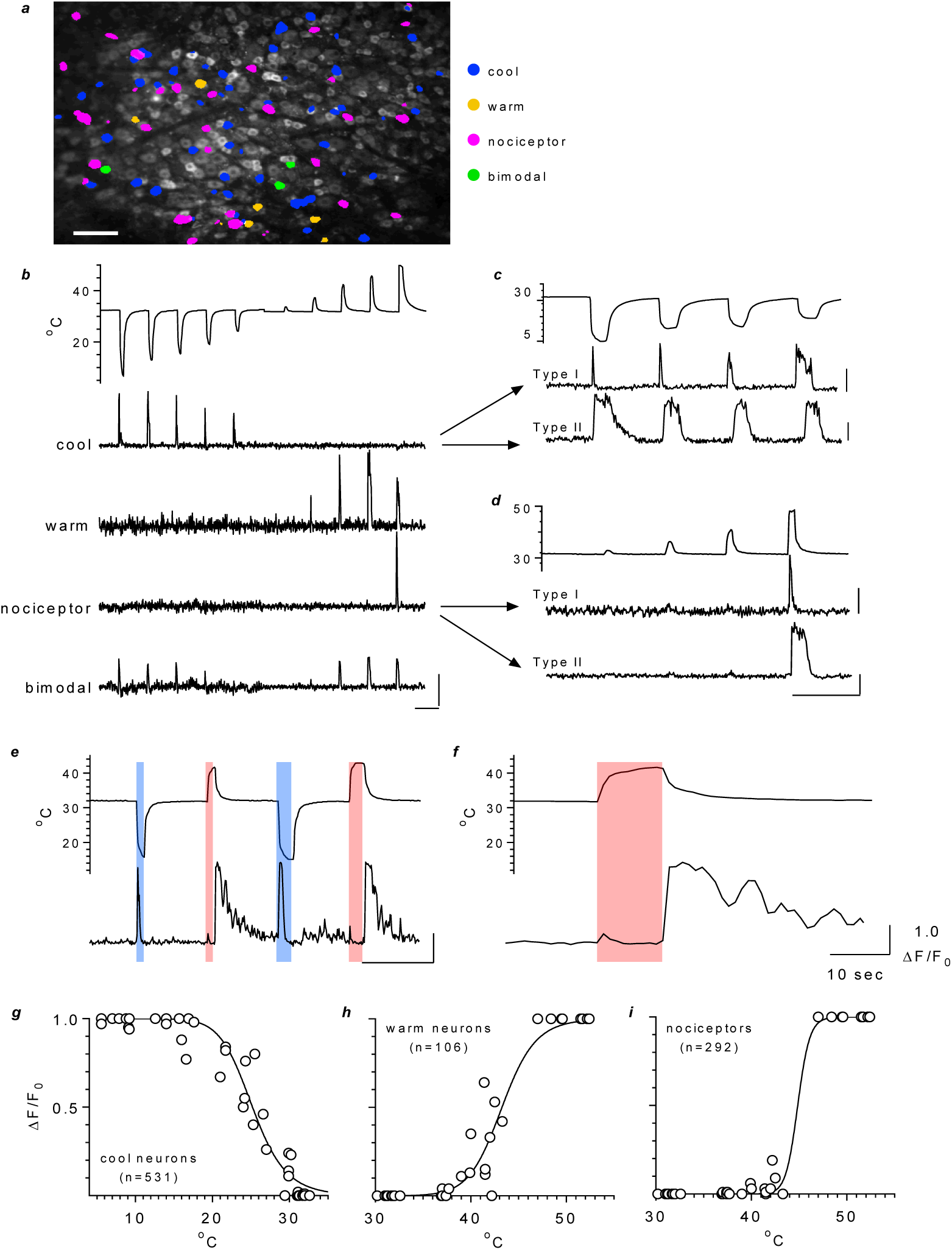
Thermosensitive trigeminal ganglion neurons in mice respond to cooling and heating the oral mucosa. Artificial saliva cooled or heated to different temperatures and flushed through the oral cavity for 10 sec elicited responses in a subpopulation of neurons. ***a***, enlarged view of the dorsal surface of area V3 of trigeminal ganglion, showing field of neurons expressing GCaMP3. Thermosensitive neurons (color-coded as shown on right) do not appear to be grouped according to thermal sensitivity. Calib, 100 microns. ***b***, top trace, a miniature thermocouple probe inserted in the buccal cavity measured the oral temperature changes from a resting temperature of 32°. Lower traces, concurrent responses from representative cool- and warm-sensing neurons, nociceptors, and bimodal-sensing neurons. ***c***, different experiment, displayed on an expanded time axis to illustrate rapidly adapting (Type I, middle trace) and slowly adapting (Type II, bottom trace) cool-sensing trigeminal neurons. Oral mucosal temperature (top trace), resting level = 32 °C. ***d***, another experiment, displayed as in c and illustrating a rapidly adapting (Type I, middle trace) and a slowly adapting (Type II, bottom trace) nociceptor response to heat stimulus (top trace). ***e***, a subset of neurons responded to cool stimuli (blue bars) as well as to the *termination* of warm stimuli (red bars), that is, to a *decrease* of oral temperature after a warm stimulus. Top trace, oral temperature; bottom trace, concurrent record from a cool-sensing neuron. Calib for ***b-e***, 1 ΔF/F_0_, 100 sec. ***f***, same data as e displayed on an expanded time scale to highlight a response following warm stimulation. Calib, 1 ΔF/F_0_, 10 sec. ***g,h,i,*** temperature-response relations for cool- and warm-sensing neurons and nociceptors (n=10 mice in each plot, number of neurons as specified). Each symbol shows the mean response at the specified temperature (n=3 to 79 neurons per point). A sigmoidal best-fit line (GraphPad Prism v.6) is drawn through the points.

### Cool- and cooling-responsive ganglion neurons

Neurons responding to thermal stimuli below 32° (“cool neurons”) comprised the largest population of thermosensitive neurons (531/1090, 49%; or ∼5% of 11,700 V3 trigeminal neurons imaged). Some cool neurons responded only transiently, especially to the lowest temperatures tested (Fig. 1c). That is, the responses rapidly adapted to the changing temperature. These neurons are similar to the Type I cold neurons described by Yarmolinsky et al. (2016). An equal proportion of cool neurons did not adapt as rapidly to decreasing temperature (Fig. 1d), similar to the Type II neurons in Yarmolinsky et al. (2016). A small proportion of cool-sensitive neurons responded only at cold extremes (<10 °C), were rapidly adapting across the full cold range, or had response patterns similar to Type III neurons (Yarmolinsky et al., 2016) (data not shown).

We also observed a novel feature of trigeminal ganglion neuron cooling responses. Namely, some cool-responsive neurons responded to *decreasing* oral temperatures (i.e. -ΔC°/Δt, “cooling”) even if the oral mucosa had first been warmed. That is, these neurons responded to the cooling phase of a warm stimulus—the return to baseline temperature (Fig. 1e,f). Similar responses have been recorded from cutaneous cool thermoreceptors (Wang et al., 2018). Cooling-sensitive responses were distributed among Types I and II cool neurons, with a greater proportion found in Type I (rapidly adapting) cool neurons (88 and 34 neurons, respectively).

Interestingly, the averaged response of all cool neurons shows minimal temperature-dependence at oral temperatures below 15∼18°(Fig. 1g). However, when examined in greater detail (below), different populations of cool neurons show distinctly different temperature-response relations across the range from 5 to 32°.

### Warm-responsive

In agreement with others, we found TG neurons that responded to thermal stimulation between 32° and 45° were the smallest group of thermosensitive neurons (9%, 106/1090). Response amplitudes in these neurons were temperature-dependent, consistent with encoding temperature intensity, and reached a maximum at ∼45° (Fig. 1b), resulting in sigmoidal population responses (Fig. 1h).

### Nociceptors

29% of oral thermoresponsive trigeminal ganglion neurons (292/1090 neurons) did not respond until the oral temperature exceeded 43-45° (Fig. 1b). As noted above, neurons with this high temperature threshold were presumed to be nociceptors. Similar to Types I and II cool-responding neurons, nociceptors were rapidly- or slowly-adapting, respectively (Fig. 1d).

### Bimodal-responsive

13% of oral thermosensitive ganglion neurons (154/1090 neurons) responded to both cool and elevated temperatures (Fig. 1a). Bimodal neurons generally showed temperature-dependent responses with increasing response amplitudes towards both temperature extremes.

### High- and low-threshold cool neurons

Another method to categorize cool neurons is based on temperature threshold. As shown by others (Bautista et al., 2007; Madrid et al., 2009), we observed that some cool thermosensing TG neurons responded to small decreases in oral temperature (“Low threshold cool-sensing neurons”, or LT) whereas other neurons did not respond until much colder temperatures were reached (“High threshold cool-sensing neurons”, or HT). We measured the approximate thresholds for 499 cool neurons (10 mice) where temperature-response relations were collected (Fig. 2a). Neurons that began responding when the oral temperature was decreased from 32° to 20° were labeled LT cool-sensing and comprised 78% (390/499) of the population. The remaining neurons that had thresholds below 20° were identified as HT cool-sensing neurons. These proportions are consistent with data from mouse trigeminal ganglion thermoreceptor neurons recorded *in vitro* (Madrid et al., 2009). Temperature-response relations differed between LT and HT cool thermosensitive neurons (Fig. 2b,c). For example, some LT cool thermosensors appeared to be “tuned” to specific temperatures; their responses dropped off at temperatures above and below an optimal value (Fig. 2c). Neurons similarly tuned to specific temperatures were also reported for cool thermoreceptors innervating the mouse hind paw (Wang et al., 2018).

**Figure 2.**
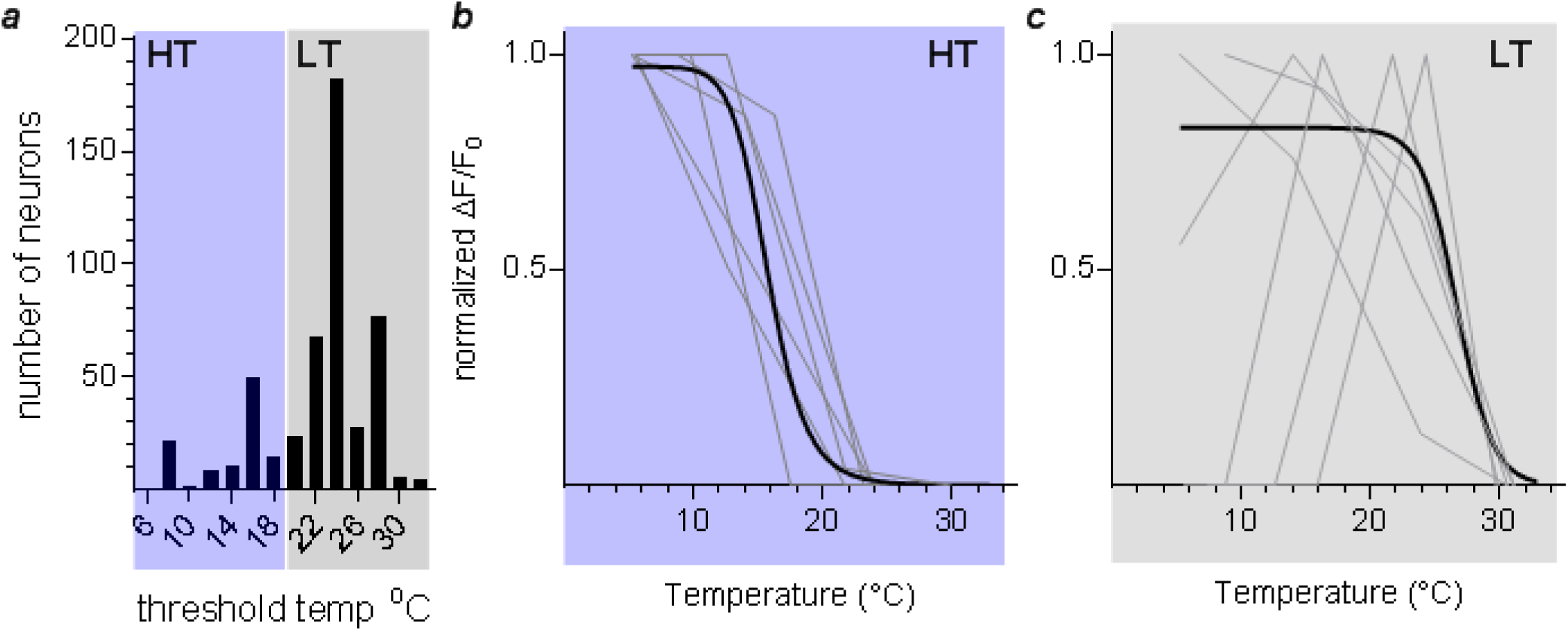
Thermosensensitive responses of high-threshold (HT) and low-threshold (LT) cool-sensing trigeminal ganglion neurons. ***a***, distribution of temperature thresholds for neurons responding to oral cooling from 32 °C. Grey shading, the majority of cool neurons (78%) responded to small temperature decreases (Δt ∼10 °C) from resting oral temperature (32 °C) (390/499 neurons, 10 mice). These are defined as low-threshold (LT) cool neurons. Blue shading, the remaining 109 thermosensitive neurons (22%) did not respond until oral temperature reached 18° or below. These are defined as high-threshold (HT) cool neurons. ***b***, temperature-response relations for HT cool neurons (n=109). Gray lines show responses for 6 representative neurons. Dark line is best-fit sigmoidal curve (GraphPad Prism) drawn for all 109 neurons. ***c***, temperature-response relations for 390 LT cool neurons. As in b, gray lines show data for 6 representative neurons and dark line is best bit sigmoidal curve for all neurons. Note that some LT cool respond best (are “tuned”) to a *specific* temperature between 6° and 32°, a phenomenon previously seen in recordings from cutaneous and trigeminal cool thermosensors (Bautista et al., 2007; Madrid et al., 2006; Wang et al., 2018).

### Population and size distributions

Figure 3a summarizes the distribution of the different types of oral thermosensitive trigeminal neurons. Oral thermosensitive neurons were significantly smaller than the average size (23.4±6.1 µm) of V3 trigeminal neurons (Fig. 3b), consistent with dorsal root ganglia thermoreceptor neuron (Caterina et al., 1997; Mizushima et al., 2006; Facer et al., 2007). There were no significant differences in size among the thermoreceptor groups (p=0.27 to 0.96, ANOVA).

**Figure 3.**
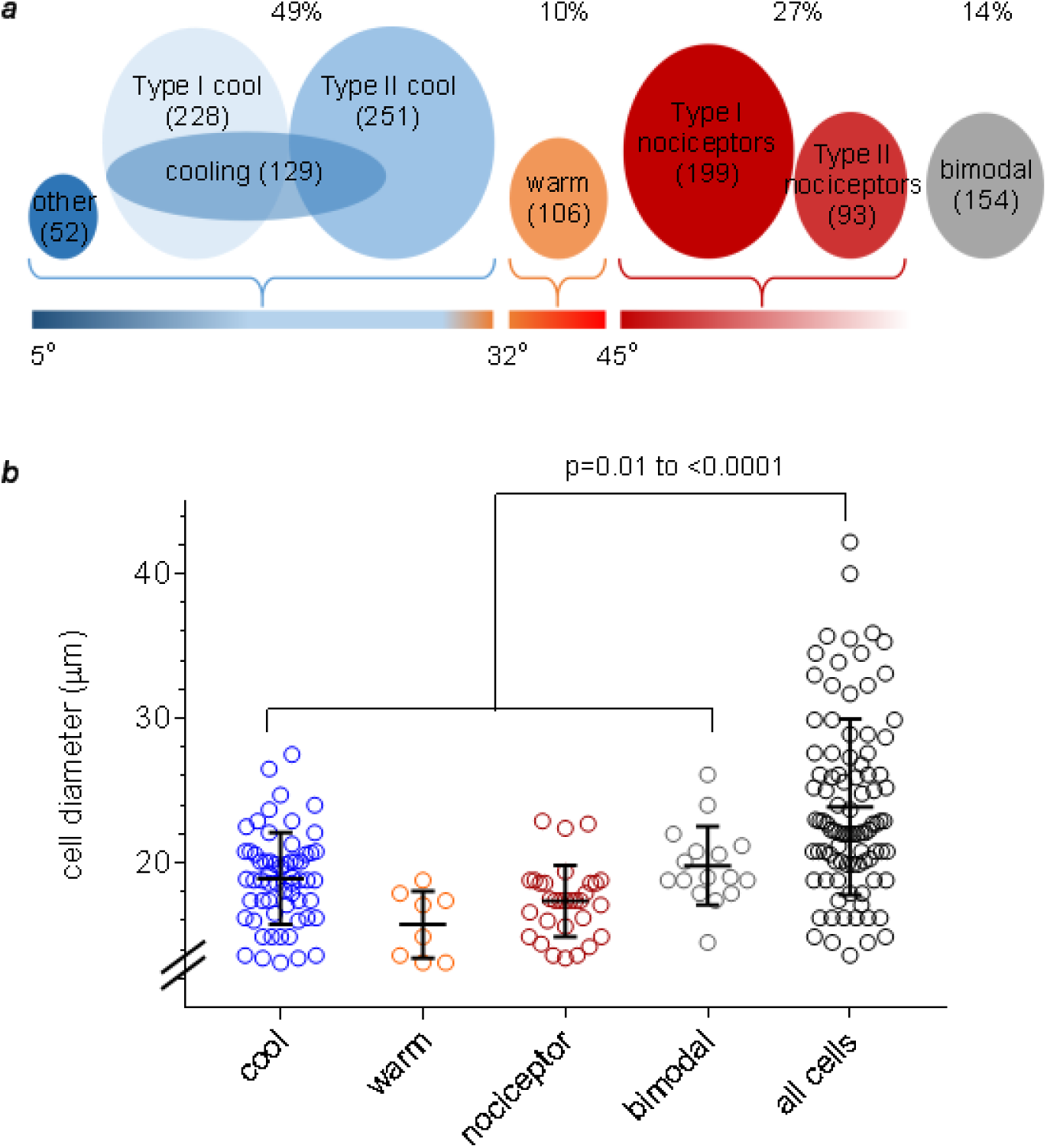
Distribution and sizes of orothermosensitive neurons in the V3 region of the trigeminal ganglion. Neurons that responded to oral temperatures were categorized into four main groups: cool responding (<32 °C), warm responding (32°to 45°), nociceptors (>45°) and bimodal. ***a***, Venn diagram illustrating the distribution of these neurons. ***b***, distribution of diameters of neurons of each category. Error bars show mean ± s.d.

### Naturally occurring pungent and cooling compounds elicit responses in trigeminal ganglion neurons

Many herbs and plants used as food condiments or in oral hygiene products, including chili peppers, mustard, and mint, generate oral thermal sensations. To investigate sensory afferent responses representing these sensations, we flushed the oral cavity of mice with the primary active ingredients of these plants— capsaicin, allyl isothiocyanate (AITC), and menthol, respectively— at concentrations found in spices or other commercial products. Capsaicin, the pungent compound in hot chili peppers, was used at 30-100 µM. Capsaicin has been reported to be ∼1000 µM in Tabasco® sauce (Betts, 1999; Kachoosangi et al., 2008). AITC evokes a pungent sensation and varies from 0.3 to 1.3 mg/g in commercial mustard (Pelosi et al., 2014) was applied at 10 mM (∼1 mg/ml). Menthol, used as a cooling agent in mouthwashes and toothpastes, was applied at 3 mM (e.g., Listerine® = 2.7 mM (Datasheet, 2008).

We observed 7% of trigeminal neurons (521/7400 neurons; 7 mice) responded to oral stimulation with capsaicin, AITC, or menthol (Fig. 4). All three compounds elicited robust responses and each compound showed a distinct response pattern (Fig. 4a). At these concentrations, the majority of responding trigeminal neurons (75%, 392/521 neurons) were activated specifically by only one of the compounds; a smaller group (22%, 115/521 neurons) responded to two compounds; and an even smaller population (3%, 14/521 neurons) responded to all three compounds (Fig 4b). Peak responses to capsaicin (1.7 ± 1.2 ΔF/F_0_) were significantly larger than those to menthol (1.1 ± 1.0 ΔF/F_0_, p<0.0001) or AITC (0.9 ± 0.6 ΔF/F_0_, p=0.0006) (Fig. 4c). Capsaicin responses also differed markedly in their time to onset and durations. Responses to capsaicin had longer latencies (97 ± 105 sec, n=58) than those to menthol (19 ± 17 sec, n=71, p<0.001) or AITC (22 ± 19 sec, n=41, p<0.0001). Capsaicin-evoked activity was significantly prolonged (180 ± 153 sec, n=58) relative to menthol (23 ± 19 sec, n=70, p<0.0001) or AITC responses (44 ± 36 sec, n=38, p<0.0001). Taken together, this likely reflects a more gradual penetration of capsaicin into and longer washout from the lingual epithelium.

**Figure 4.**
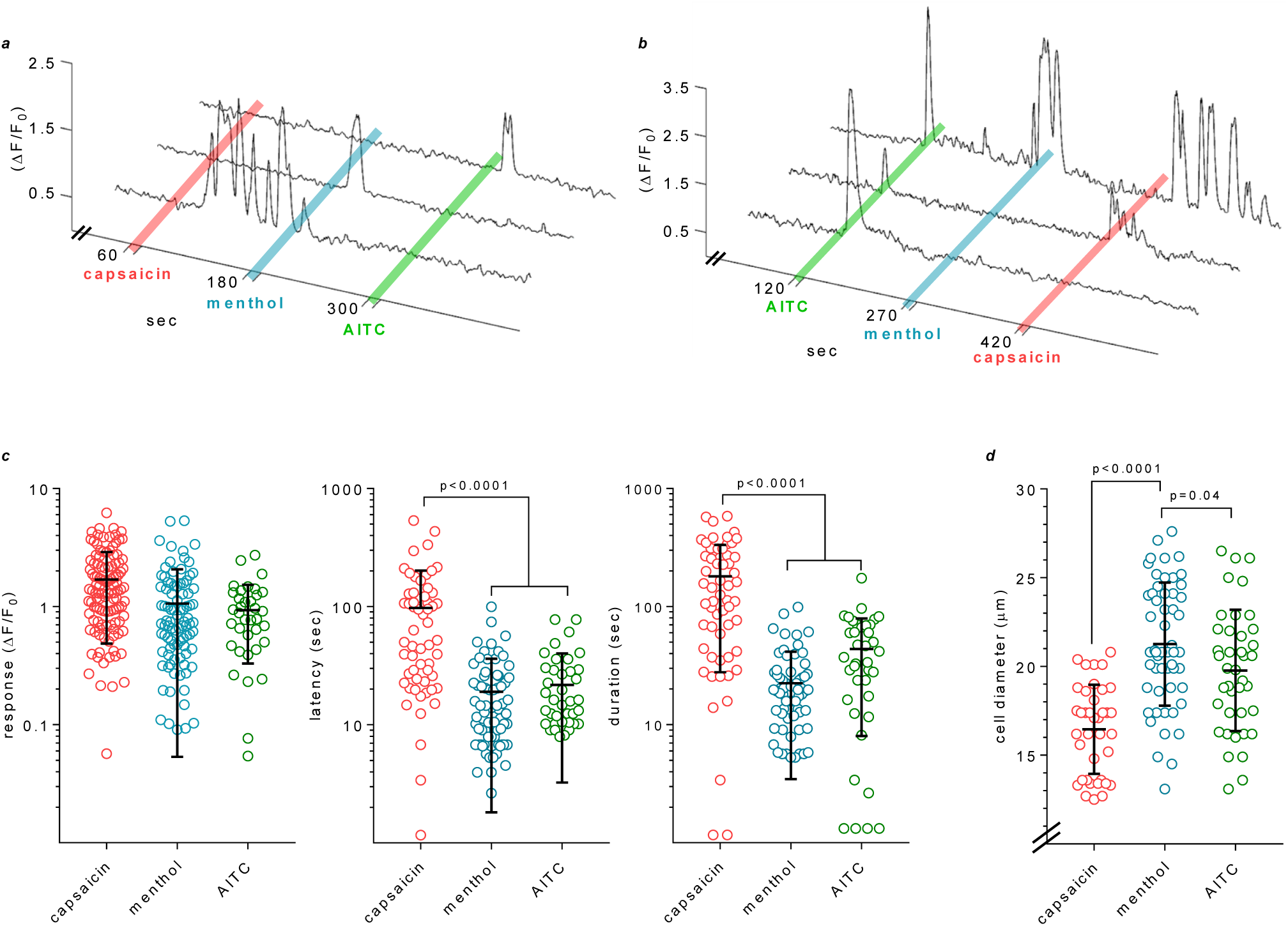
Trigeminal ganglion neurons respond to oral stimulation with capsaicin, menthol, and mustard oil (AITC). ***a,*** representative responses of 3 trigeminal ganglion neurons that respond selectively to 10 sec oral lavage with capsaicin (100 µM), menthol (3 mM), and AITC (10 mM). Notice the delayed onset and prolonged response to capsaicin, which was typical for this compound. ***b,*** 3 neurons in another experiment respond to 1, 2 or all 3 compounds. ***c***, amplitudes (ΔF/F_0_), latencies, and durations of responses to capsaicin, menthol, or AITC. Note, ordinates are logarithmic scales to display the variability. ***d***, diameters of trigeminal neurons that responded exclusively to capsaicin, menthol, or AITC (as in *a*, above). Error bars show means ± s.d.

Neurons that responded to menthol and AITC were of similar sizes (Fig. 4d), having diameters of medium-sized sensory neurons (21.3 ± 3.5 µm, n=54 and 19.8 ± 3.4 µm, n=40, respectively). Capsaicin-responsive neurons were significantly smaller (16.5 ± 2.5 µm, n=38; p<0.0001). As a generality, neurons responding to menthol, AITC or capsaicin were significantly smaller than the average size of trigeminal ganglion neurons (23.4 ± 6.1 µm, n=94; p <0.0001).

### Trigeminal thermoreceptors respond to menthol, AITC, and capsaicin

Given that the compounds tested stimulate oral perceptions of warmth or cool, we next assessed whether there was an interaction between the thermosensitivity of a trigeminal neuron and its sensitivity to capsaicin, AITC, or menthol. For these experiments (416 neurons; 3 mice), an initial test of stimuli spanning the entire temperature range (∼5 to 52 °C, see Fig. 1) was conducted to identify and classify trigeminal thermoreceptors into the 4 groups: cool neurons, warm neurons, bimodal thermoreceptors and nociceptors. Following this, menthol, AITC, and capsaicin were applied sequentially, with their order counterbalanced between experiments. Across the three experiments, nearly half the thermosensitive trigeminal ganglion neurons (204/416) responded to menthol, AITC, capsaicin, or a combination of these compounds. As was the case for thermosensitive neurons, there was no spatial mapping of capsaicin-, AITC-, or menthol-sensitive neurons in the V3 area of the trigeminal ganglion. Figure 5a is a Venn diagram representing the incidence of responses to menthol, AITC, and capsaicin among trigeminal thermoreceptors. The majority of neurons that responded to capsaicin, AITC, or menthol were also thermosensitive (204/270).

**Figure 5.**
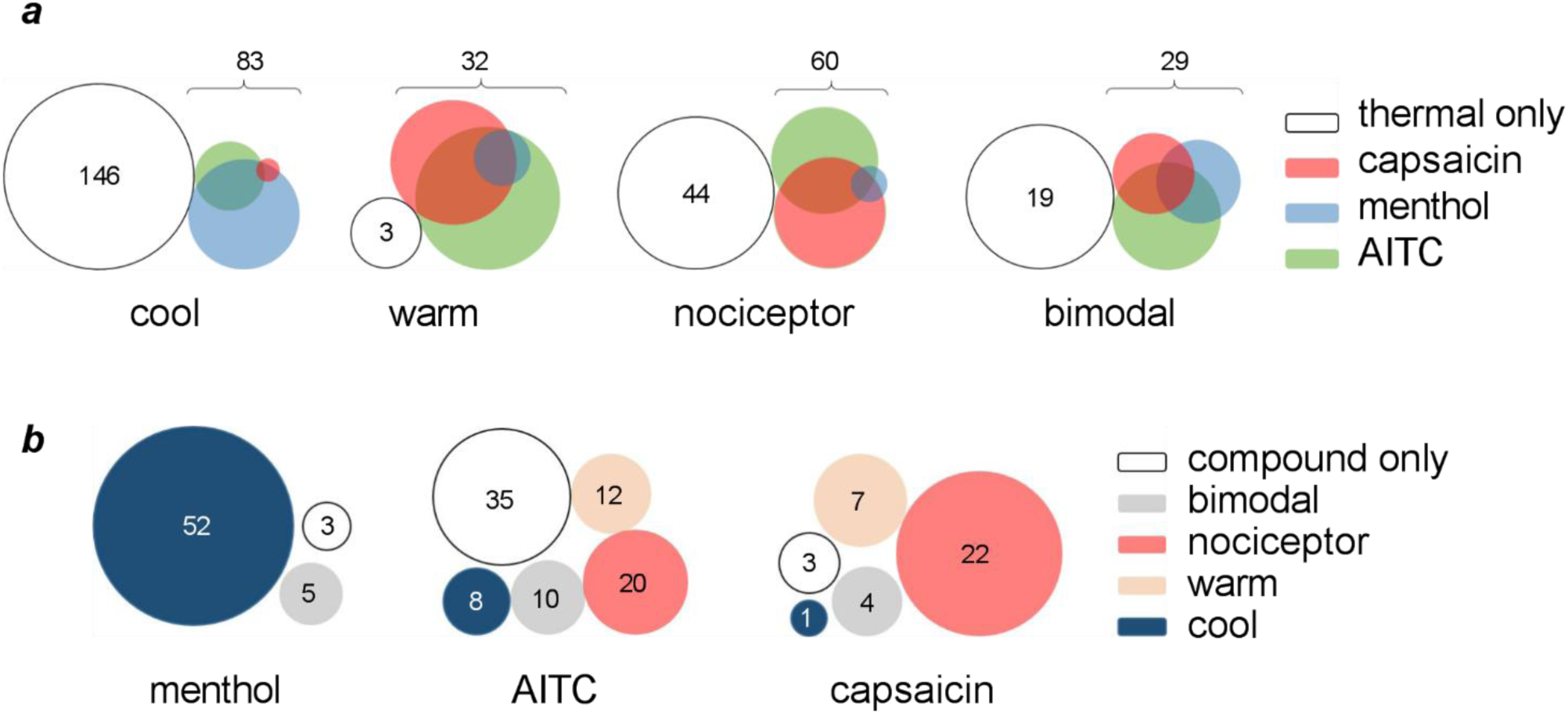
Thermosensing trigeminal ganglion neurons also respond to oral stimulation with capsaicin, menthol, or AITC. ***a***, Venn diagrams illustrating the distribution of thermosensing trigeminal ganglion neurons and their sensitivity to capsaicin, menthol, and/or AITC (3 mice, 416 neurons). The sizes of the Venn diagrams have been normalized across thermosensing categories to illustrate the proportional distributions (e.g., there are far fewer warm neurons in the sample). Note that even though certain classes of thermosensing neurons respond predominantly to one of the compounds (e.g., cool neurons predominantly respond to menthol), there nonetheless is considerable overlap in chemosensitivity. ***b.*** Conversely, this Venn diagram illustrates how neurons that responded exclusively to menthol, AITC or capsaicin (3 mice) were distributed among trigeminal ganglion neurons probed with thermal stimuli. The category “compound only” indicates neurons responding only to chemical, but not to thermal stimulation.

Thermosensitive neurons had characteristic responses to the chemesthetic compounds. Namely, the greatest proportion of cool neurons responded to menthol, either alone or in combination with capsaicin and/or AITC (74/83; 89%). Warm-sensitive neurons had a greater incidence of capsaicin-(19/32; 59%) and AITC-sensitivity (25/32; 78%). Similarly, trigeminal nociceptors were most sensitive to capsaicin (39/60; 65%) and AITC (36/60; 60%). When considering the neurons that responded exclusively to one of the three tested compounds (Fig. 5b), menthol-sensitivity (60 neurons) is overrepresented in cool-sensitive neurons and capsaicin-sensitivity (37 neurons) in nociceptors. Interestingly, a large portion (41%) of the neurons responding exclusively to AITC (85 neurons) were thermally insensitive. Thermally insensitive neurons with responses to menthol or capsaicin were less common (3/60; 5% and 3/37; 8%, respectively). In short, though there was a tendency for cool neurons to respond to menthol and vice versa, and a tendency for warm neurons and nociceptors to respond to capsaicin and vice versa, these associations were far from absolute.

Studies indicate that there are relative differences in TRPM8 and TRPA1 channel expression in low-threshold versus high-threshold trigeminal cool sensitive neurons (Madrid et al., 2009; Teichert et al., 2014). Thus, we tested whether low- and high-threshold oral cold thermosensitive trigeminal ganglion neurons differed in their responses to menthol, AITC, or capsaicin. There was no significant difference between LT and HT cool-neurons regarding their activation by AITC or capsaicin. Specifically, AITC stimulated 31% (11/35) HT versus 38% (42/110) LT cool neurons; capsaicin activated 15% (7/43) HT versus 16% (21/140) LT cool neurons. In contrast, a significantly greater fraction of LT versus HT cool trigeminal ganglion neurons was stimulated by menthol: 44% (87/199) versus 19% (3/16). (p=0.001, Fisher’s Exact Test, 2 tailed). The predominant sensitivity of LT cool neurons to menthol is consistent with findings on mouse trigeminal and dorsal root ganglion neurons *in vitro* (Madrid et al., 2009; Teichert et al., 2014).

### Capsaicin causes thermal sensitization

Finding that menthol, AITC, and capsaicin all stimulate thermoreceptors, we next explored whether these compounds exerted any lasting effects on thermosensitivity in the oral cavity. Spices containing capsaicin such as chili peppers notoriously enhance and prolong a sense of heat in foods (Green, 1986b).

To begin this series of experiments, we flushed the oral cavity with warm artificial saliva (41 to 42 °C) twice at 90 sec intervals followed by a 10 sec oral lavage with 100 µM capsaicin. After capsaicin, 41-42° stimuli were re-applied. Capsaicin markedly increased the numbers of neurons responding (Fig. 6). When tested in triplicate (3 mice), we observed a >2.5 fold-increase in the number of neurons responding to 41-42 °C stimulation after capsaicin and a ∼6.5-fold increase in the responses (ΔF/F_0_) (Fig. 6b,c).

**Figure 6.**
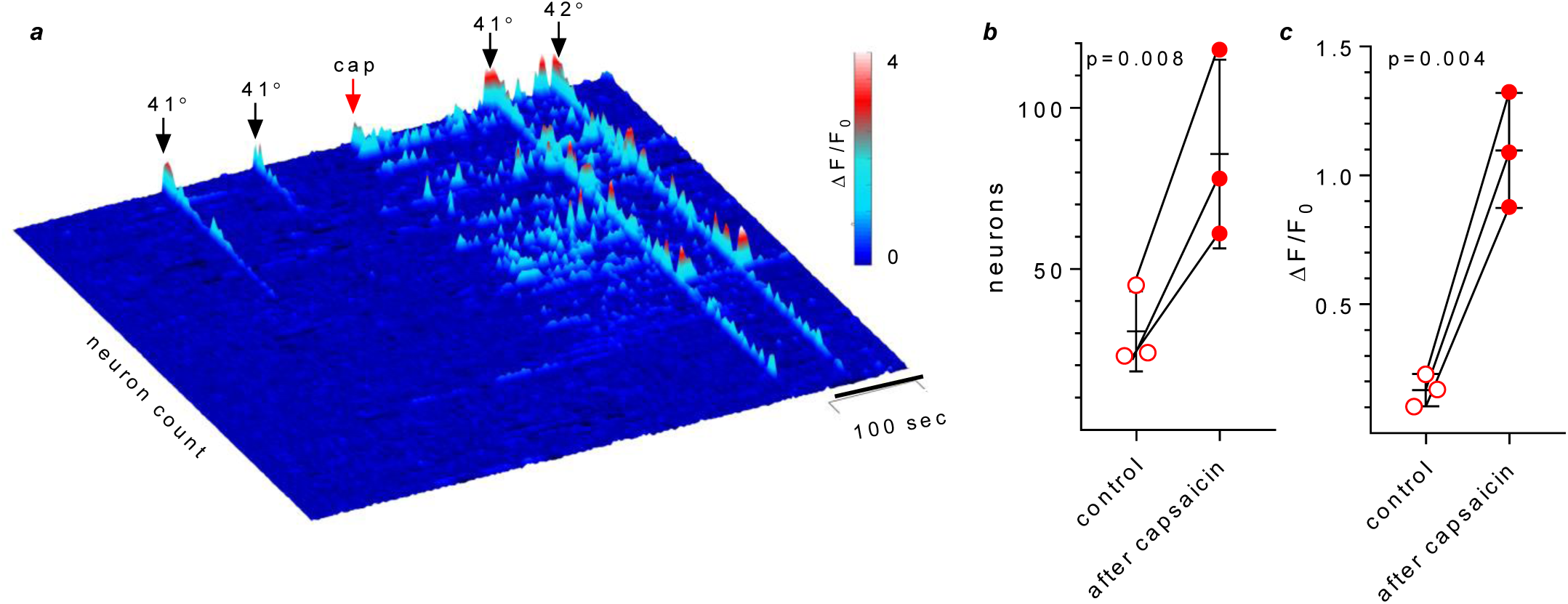
Capsaicin sensitizes neurons innervating oral mucosa to warm stimuli. ***a***, heat map showing responses from trigeminal neurons (n=107) to warm (41-42 °C) artificial saliva (AS), applied twice for 10 sec each, before and following a 10 sec lavage with 100 µM capsaicin (cap, 32 °C). Note the increase in number of warm-responding neurons and amplitudes of responses after capsaicin. ***b***,***c,*** summary of data from 3 mice (257 neurons) as in ***a***, presented as before/after plots. Each symbol shows the data from one animal, *viz*. the mean of 2 presentations of artificial saliva (41-42°) before and after capsaicin. Error bars show means ± s.d.

### Capsaicin causes warm sensitization and thermal hyperalgesia

To identify trigeminal thermoreceptors sensitized by capsaicin lavage, we refined the above experiments by testing a complete temperature range (∼5 to 52 °C), before and after oral lavage with 100 µM capsaicin. This enabled us to identify and track cool neurons, warm neurons, and nociceptors prior to and following the application of capsaicin. Parenthetically, stimulating with nociceptive hot solutions for 15 sec or longer can produce changes in oral thermoreceptors due to mucosal inflammation (see Yarmolinsky et al., 2016). Thus, we identified nociceptors by briefly (10 sec) applying artificial saliva at a range of temperatures only up to 52°, followed immediately by a cold rinse (10 to 15°) and a ∼10 minute rest. This protocol induced minimal to no mucosal inflammation, as confirmed by re-testing temperature-response relations and observing no significant difference from the initial trial. After recording two temperature-response relations (controls), we applied capsaicin and repeated the temperature-response series. The same protocol was followed in triplicate experiments (3 mice). The results unambiguously showed that oral lavage with capsaicin causes trigeminal nociceptors, which normally have a threshold >43 to 45 °C now to respond to innocuous warm temperatures (35-43 °C) (Fig. 7). Responses before and after capsaicin in a representative nociceptor are shown in Fig. 7a. Temperature-response curves for identified nociceptors obtained before and after capsaicin show a pronounced left-shift (EC_50_ 45.0 vs 39.5 °C, respectively), indicating that the threshold for nociceptors shifted to lower temperatures (i.e., thermal hyperalgesia) (Fig. 7b). This left-shift in temperature-response relations after capsaicin resulted in 56 % (45/81) of the previously identified nociceptors now responding to the warm range (32-45 °C). That is, the “novel” responses in Fig. 6 are in fact generated by sensitized nociceptors. Similarly, previously identified warm neurons also had lower thermal thresholds after capsaicin (EC50 41.5 vs 36.5, respectively). They, too, are more responsive to mild warm temperatures (Fig. 7c). Cool neurons were unaffected by capsaicin at this concentration. Bimodal neurons showed sensitization limited to warm temperatures with increased response amplitudes but with an unchanged incidence (data not shown). In short, thermal hyperalgesia produced by capsaicin can be explained by sensitization of nociceptors and warm neurons (and possibly bimodal thermoreceptors).

**Figure 7.**
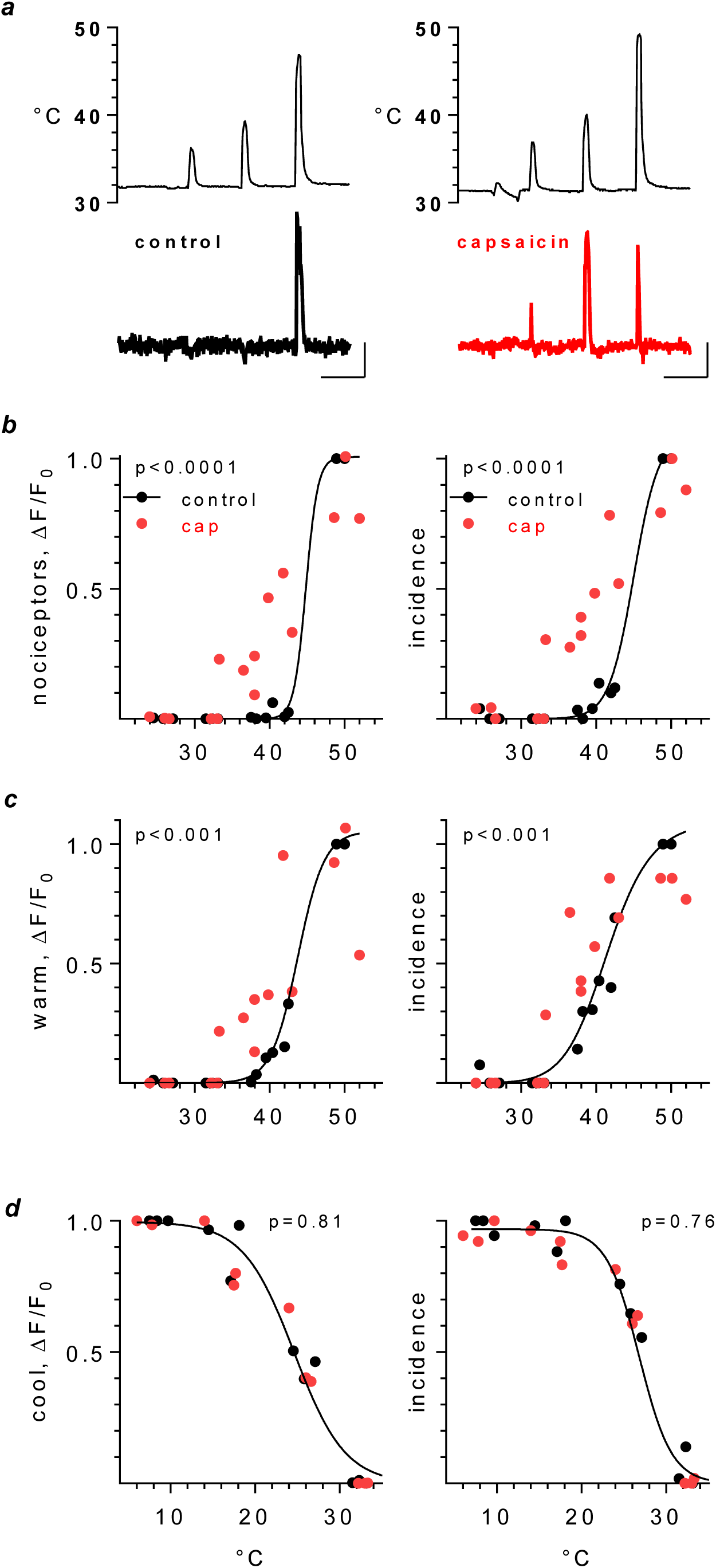
Pretreatment with capsaicin sensitizes nociceptors and warm-sensing trigeminal ganglion neurons. ***a***, Response of a representative nociceptor to thermal stimulation before (left) and after (right) a 10 sec lavage with 100 µM capsaicin. Note that after capsaicin this nociceptor began to respond to warm (<45°) thermal stimulation. Upper traces, temperature of oral mucosa. Calib, 100 sec, 0.5 ΔF/F_0_. ***b,c,d,*** temperature-response relations for nociceptors, warm, and cool trigeminal ganglion neurons (3 mice, 81, 28 and 150 neurons, respectively), followed before (black symbols) and after (red symbols) oral capsaicin lavage. Each point shows data from a single mouse. For each row of temperature-response relations, the left plots show response amplitudes (mean ΔF/F_0_) and the middle plots show response incidences. The data for each mouse have been normalized to the maximum values obtained before capsaicin. Solid line is best-fit sigmoidal curve (GraphPad Prism) for data *before* capsaicin. p values show significance of difference between before/after data in each plot.

### Menthol and mustard oil alter responses of oral thermoreceptors

The effects of menthol and AITC on the temperature-evoked responses of trigeminal thermoreceptors were more ambiguous and less striking than those of capsaicin. For example, menthol (3 mM) had no clear effects on the amplitudes of nociceptors and warm neurons (Fig. 8a,b, 3 mice), but significantly reduced the amplitudes of responses of cool neurons (Fig. 8c). At the same time, menthol had no significant effect on the incidence of warm- or cool-responding neurons, but somewhat reduced the incidence of nociceptors (Fig. 8a). Menthol did not appear to shift the threshold for cool neurons (Fig. 8c), unlike the effects of menthol applied directly onto isolated trigeminal ganglion neurons *in vitro* (McKemy et al., 2002). Conversely, after 10 mM AITC, 41% (24/59) of previously identified nociceptors now responded in the warm range; cool neurons were not affected (Fig. 9, 3 mice). AITC increased the response amplitudes of warm-responding neurons and nociceptors (though the latter could not be verified statistically). Lastly, AITC seems to reduce the amplitude of responses in cool neurons but the significance of this effect could not be quantified.

**Figure 8.**
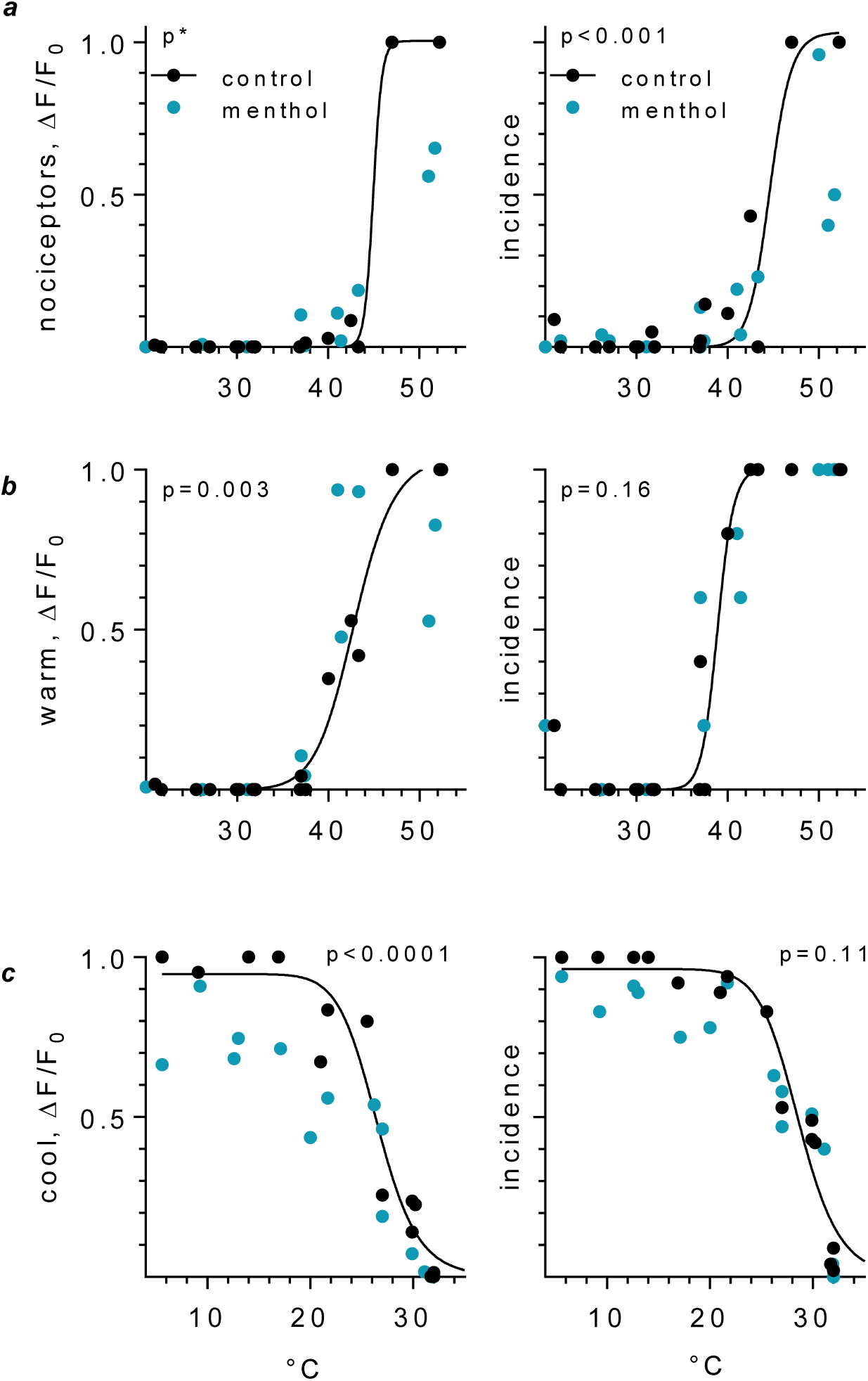
Pretreatment with menthol affects oral thermosensory responses in trigeminal ganglion neurons. ***a,b,c,*** temperature-response relations for nociceptors, warm, and cool trigeminal ganglion neurons (3 mice, 99, 16 and 122 neurons, respectively), followed before (black symbols) and after (teal symbols) oral menthol (3 mM) lavage, organized and displayed as in Fig. 7 b-d. The most pronounced effect of menthol was to decrease the amplitude of cool neuron responses, *c*, and, to a lesser extent, reduce the incidence of nociceptors, *a*. Curves drawn as in Fig. 7; p*, data cannot be fit to a simple model for statistical comparisons by GraphPad Prism (v.6).

**Figure 9.**
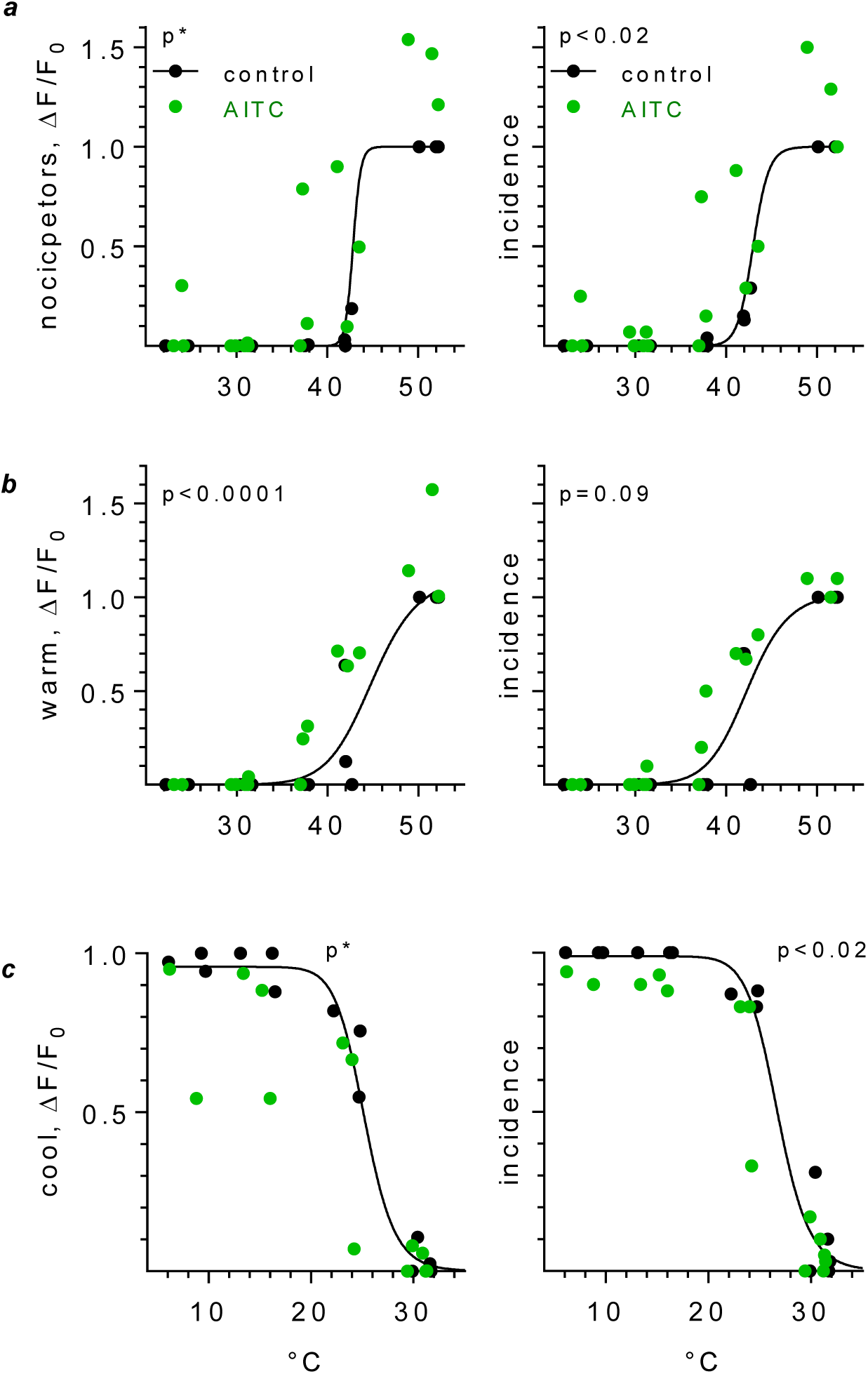
Pretreatment with AITC affects oral thermosensory responses in trigeminal ganglion neurons. ***a,b,c,*** as in Figure 7 *b-d*, temperature-response relations for nociceptors, warm, and cool trigeminal ganglion neurons (3 mice, 59, 26 and 139 neurons, respectively), followed before (black symbols) and after (green symbols) oral AITC (10 mM) lavage, organized and displayed as in Fig. 7 b-d. AITC increases the amplitude of warm neuron responses, *b*, and to a lesser extent, increases the incidence of nociceptors, *a*, and decreases that of cool neurons, *c*. Curves drawn as in Fig. 7; p*, data cannot be fit to a simple model for statistical comparisons by GraphPad Prism (v.6).

## Discussion

Our findings show that thermal stimulation of the oral cavity, particularly cool, is effective in activating large numbers of trigeminal thermosensory ganglion neurons. Some neurons respond to warm and cool temperatures alike whereas others are tuned to a specific temperature (Figs. 1, 2). Our results are consistent with thermal orosensations being generated by population coding by the trigeminal ganglion (see below). Moreover, certain spices and natural compounds found in foods and oral hygiene products, namely capsaicin, mustard oil, and menthol, mimic and interact with these orosensory thermal stimuli.

### Cool and warm thermosensing

In the V3 region of the trigeminal ganglion there is a predominance of cool-versus warm-responding sensory neurons. This predominance of cool-sensing afferent innervation was observed since early recordings from the lingual nerve and later (Zotterman, 1936; Yamashita et al., 1964; Wang et al., 1993). Based on the large population of cool-sensing trigeminal neurons innervating the oral cavity, one might predict that mice detect and are more sensitive to cool versus warm stimulation of the tongue, palate, and cheeks, but we are unaware of any such studies. The closest parallel is that mice appear to be more sensitive to *cutaneous* cooling than to warming (Montesinos et al., 2017) despite there being a predominance of warm-over cool-sensing C fibers innervating the skin (Jankowski et al., 2017). Behavioral preference testing indicate that rats detect and discriminate drinking water in the same temperature ranges tested in our study, though threshold detections were not studied (Kapatos & Gold, 1972; Deaux, 1973; Smith et al., 2010; Torregrossa et al., 2012). Human subjects either (a) have difficulty identifying small temperature changes when applied focally to the tongue (Niissalo et al., 2003), or (b) show no difference in detection thresholds for increased vs decreased temperatures (Granot & Nagler, 2005; Kalantzis et al., 2007), or (c) are more sensitive to cooling than to warming (Green, 1984; Renton et al., 2003). When whole-mouth thermosensing was assayed with temperature-controlled liquids, subjects appeared to be more sensitive to warmth (Green, 1986a). For comparison, Figure 10 shows whole mouth intensity ratings for cooling and warming from human subjects (data from Green (1986a)) plotted on the same temperature scale with mouse V3 trigeminal thermosensing neurons (data from Fig. 1f,g), indicating approximate parallels between mice and humans.

**Figure 10.**
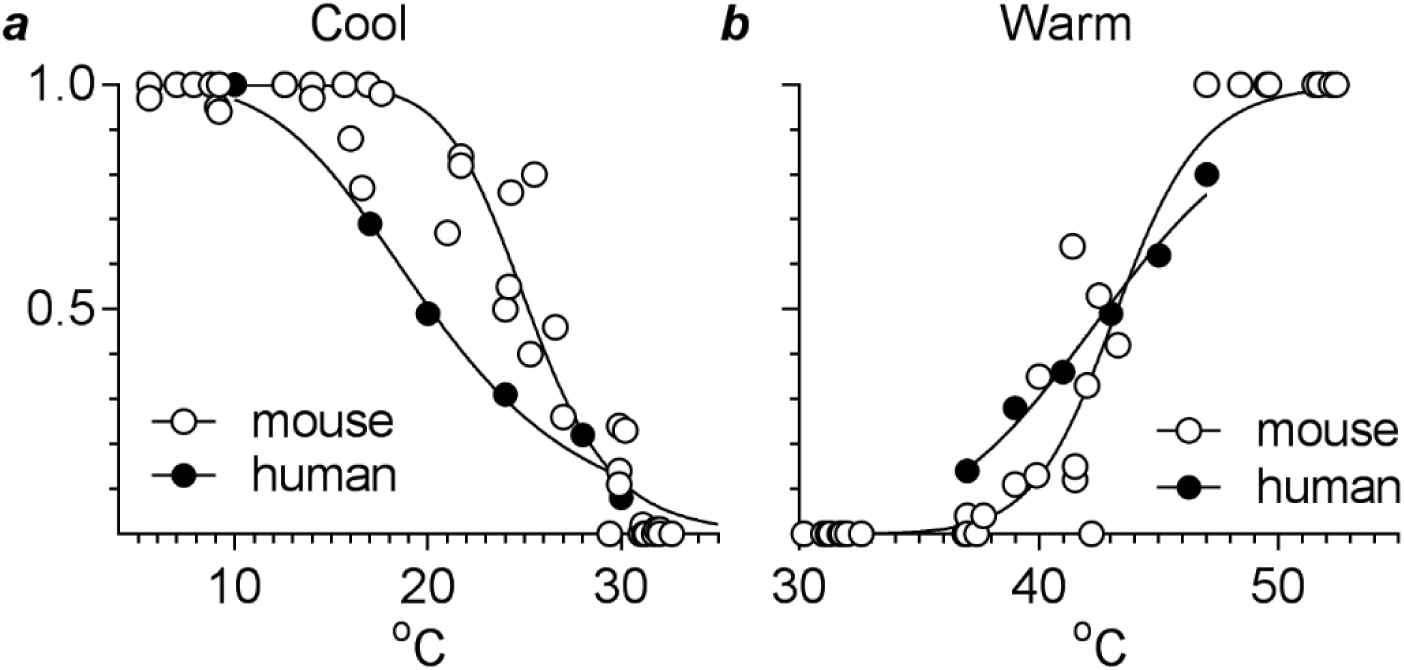
Human psychophysical evaluation of oral temperatures compared with mouse trigeminal thermosensory neuron stimulation. ***a,*** intensity rating for cool liquids by human subjects (●) overlaid onto cool temperature-response data from mouse trigeminal orothermosensory neurons (○, data from Fig. 1g). Y axis shows psychophysical intensity ratings normalized to the maximum cold intensity reported, from Green (1986a). ***b,*** similar comparison for warm temperatures, normalized as in ***a*** to maximum intensity reported for cold (Green, 1986a) and including mouse data from Fig. 1h). Symbols as in ***a***.

### Responses to capsaicin, menthol, or mustard oil and their modulation of thermosensitivity

Separate but overlapping populations of mouse trigeminal ganglion sensory neurons were activated by oral stimulation with capsaicin, mustard oil (AITC), or menthol. We intentionally applied capsaicin (30-100 µM), AITC (10 mM), and menthol (3 mM) at concentrations found in commonly available foods, spices, and commercial products or used in human psychophysical and animal taste behavior studies. For instance, concentrations of capsaicin up to 3 mM have been used in human psychophysical experiments, but subjects report that lower concentrations (10-20 µM) are “hot” or “burning”, though not specifically “painful” (Karrer & Bartoshuk, 1991; Kapaun & Dando, 2017). Mice avoid drinking aqueous solutions of capsaicin above 1.65 µM but do not completely reject even 330 µM capsaicin solutions (Simons et al., 2001). Mice reduce drinking of menthol solutions above 640 µM but still consume menthol at twice that concentration (Fan et al., 2016). We are unaware of data on mice consuming AITC solutions. In sum, the concentrations of oral stimuli in the present study are irritating to but not unreasonably high for human subjects and, with the exception of AITC, have been shown to be salient and not completely rejected by mice.

In addition to activating trigeminal sensory neurons, each of the above compounds altered thermosensing by the oral mucosa. Capsaicin and AITC sensitize nociceptors and warm thermoreceptors such that they respond more strongly to temperatures lower than they would in the absence of the compound. Most striking was the ability of capsaicin to sensitize nociceptors, though warm thermoreceptors were also affected. Under control conditions, thermal nociceptors were not activated by oral stimuli below ∼43°-45°. Following capsaicin lavage of the oral mucosa nociceptors were strongly stimulated merely by warm (innocuous) temperatures, i.e. > 32° (resting oral temperature). Our data resemble hyperalgesia to warm and hot stimuli that follows thermal damage (or application of capsaicin) to the skin (Bessou & Perl, 1969; Carpenter & Lynn, 1981; Culp et al., 1989). At the concentrations used in our study, cool-sensing neurons were unaffected by capsaicin.

It is well established that capsaicin, menthol, and AITC have marked effects on subpopulations of sensory ganglion neurons (Roper, 2014; Vriens et al., 2014; Simon and Gutierrez, 2017). Moreover, the ability of these chemesthetic compounds, at least menthol and capsaicin, to modulate oral thermosensing has been well studied. For instance, capsaicin enhances perceptions of warmth of solutions sipped and held in the mouth (Green, 1986b; Kapaun & Dando, 2017), corresponding to our findings of capsaicin-induced enhanced warm thermosensing (Fig. 6). Reports are conflicting on the ability of capsaicin to affect oral cool perception in human subjects; there are arguments for (Green, 1986b) and against (Szolcsanyi, 1977; Green, 2005) a capsaicin-induced reduction of cool sensitivity, depending on the methodology of testing. In the present report, pre-exposure to capsaicin did not affect cool-responses in mouse trigeminal neurons (Fig. 7d). Menthol is well known to stimulate cool perceptions on its own and enhance responses of isolated sensory neurons to cool temperatures (McKemy et al., 2002). Menthol’s effects on oral thermosensing in human psychophysical experiments are more complex. Subjects reported that cool perceptions are enhanced and warm sensations initially attenuated by menthol solutions (Green, 1985, 2005). To our knowledge, interactions between AITC and oral thermal sensing have not been reported. Apart from menthol’s reduction of cool-sensing trigeminal neurons (Fig. 8c), the effects of AITC and menthol on oral trigeminal thermosensors were small, although in some cases statistically significant. This is most likely due to the washout of these compounds (Fig. 4) during the thermal sensory tests. The actions of menthol and AITC on oral mucosa are not as prolonged as those of capsaicin.

It is tempting to explain the actions of capsaicin, AITC, and menthol as stemming from their actions on specific TRP channels in trigeminal nerve endings innervating the oral mucosa (Roper, 2014; Simon and Gutierrez, 2017). Indeed, in a recent study aimed at characterizing TRP channels in oral trigeminal thermoreceptors, Yarmolinsky et al., (2016) intentionally used high concentrations of those compounds (capsaicin, 5 mM; menthol, 640 mM; AITC, 126 mM) to activate and study trigeminal afferent neurons. Capsaicin strongly activates TRPV1, AITC stimulates TRPA1 and TRPV1, and menthol is an agonist for TRPM8. Although separate populations of trigeminal neurons preferentially express each of these TRP channels, there is overlap and co-expression of different combinations of TRPV1, TRPA1, and TRPM8. Notably, Nguyen et al. (2017) report co-expression of TRPV1 and TRPM8 in a small fraction of mouse trigeminal neurons. These neurons might correspond with the co-activation by capsaicin and menthol we report here for cool-sensing and bimodal thermosensing neurons (Fig. 5a). Similarly, other trigeminal neurons co-express TRPA1 and TRPV1 (Nguyen et al., 2017) which could well correspond with the overlap we observed between capsaicin- and AITC stimulation (Fig. 5a). It is noteworthy that some trigeminal ganglion neurons that were activated by cool, warm, or hot oral temperatures were not stimulated by menthol, AITC, or capsaicin. Conversely, a proportion of chemosensitive neurons, especially those responsive to AITC, were not thermosensitive.

However, in the oral mucosa there are other potential peripheral targets for capsaicin, AITC, and menthol. In addition to trigeminal nerve endings in the oral mucosa, keratinocytes in the oral epithelium may also be activated. Keratinocytes express TRP channels and activating keratinocytes stimulates afferent fibers (Lee & Caterina, 2005; Lumpkin & Caterina, 2007; Baumbauer et al., 2015). It is highly likely that capsaicin, AITC, or menthol directly stimulate epithelial neurons, which in turn release neuroactive substances such as ATP, β-endorphin, and others and thus secondarily stimulate trigeminal sensory fibers (Lumpkin & Caterina, 2007; Sondersorg et al., 2014).

Lastly, capsaicin, AITC, and menthol may have multiple actions, including effects on targets other than Trp channels. For instance, capsaicin, acting through TRPV1, increases intracellular cAMP, activates cGMP-PKG and CaMKII pathways, and inhibits voltage-gated sodium and potassium channels (Liu et al., 2001; Liu & Simon, 2003). Capsaicin also modulates potassium channels and inhibits neuronal action potentials via TRPV1-independent pathways (Pezzoli et al., 2014; Yang et al., 2014) and elicits inward Na currents in ∼1/4 of geniculate ganglion neurons (Nakamura & Bradley, 2011) despite the absence of TRPV1 expression in that ganglion (Dvoryanchikov et al., 2017). Further, topical capsaicin (and mustard oil) is well known to produce vasodilation and increased blood flow (Jancso, 1960; Jancso et al., 1967; Helme & McKernan, 1985; Roberts et al., 1992), which itself will alter the mucosal temperature and thus indirectly influence thermosensory responses (Wang et al., 1995). Parenthetically, in the present study changes in mucosal temperature were mitigated by constant perfusion with temperature-controlled artificial saliva. Regarding menthol, in addition to being a TRPM8 agonist and a cooling agent, this compound is a positive allosteric modulator of GABA_A_ receptors and inhibits neuronal excitability through TRP-independent pathways, possibly being related to menthol’s role as an anesthetic (Hall et al., 2004; Pezzoli et al., 2014). The depression of trigeminal thermoresponses, even to cool, by menthol (Fig. 8) may reflect menthol’s anesthetic characteristics.

In sum, oral sensations and modulation of thermosensitivity mediated by capsaicin, AITC, or menthol may have several origins, not merely activation of TRP channels in trigeminal afferent fibers innervating oral mucosa. This caveat underlies discussions of chemesthetic stimulation, orothermal sensitivity, and TRP channels in the oral cavity.

### Thermal coding in the trigeminal ganglion

Collectively, our findings are consistent with the notion that the oral mucosa senses thermal gradients by population coding. When taken collectively, population averages of cool and warm neurons show well-behaved sigmoidal temperature-response relations, whereas individual neurons can be tuned to a specific temperature or respond bimodally. Similar tuning can be seen in electrophysiological recordings from thermosensitive cutaneous fibers and trigeminal neurons (Bautista et al., 2007; Madrid et al., 2006). Such selective tuning of cold thermosensitive neurons is not readily explained. TRPM8 channels, which presumably underlie cool transduction, do not exhibit such selective tuning (McKemy et al., 2002). More likely is that tuning is a property of the excitability of trigeminal afferent nerve terminals (Viana, personal communication). This might explain the conflicting data regarding orosensitivity to cool and warm in human subjects; if orosensory thermal input is sensed by the combined population of afferent thermosensory trigeminal neurons, thermal sensations would be expected to be highly dependent on the subset of neurons activated, which likely varied from laboratory to laboratory. Extensive data from imaging thermal responses in dorsal root ganglion neurons that innervate the hind paws in mice came to the same conclusion, namely that cutaneous temperature is signaled by population coding (Wang et al., 2018). These conclusions support earlier interpretations of thermoreception coding (Ma, 2012).

## Additional information

### Author contributions

SCML, AFN and SDR designed research. SCML and AFN performed the research and analyzed data. SCML, AFN, and SDR wrote the paper. SCML, AFN, SAS, NC, SDR interpreted data and revised the work critically for important intellectual content.

### Funding

This work was funded by NIH grants 5R01DC014420 (NIDCD), R21DE027237 (NIDCR, NIHNCI) and by Ajinomoto Co., Inc.

#### Acknowledgements

We thank Professor Xinzhong Dong at Johns Hopkins School of Medicine for the generous contribution of transgenic Pirt-Cre and Pirt-GCaMP3 mice. We are also grateful to São Paulo Research Foundation (FAPESP) for the Research Internship Fellowship of AFN (BEPE Process: 2016-07838-8).

